# Females versus males exhibit greater brain activation in sensorimotor regions during motor imagery after stroke

**DOI:** 10.1101/2025.03.06.641936

**Authors:** Sara M. Klick, Justine R. Magnuson, Lauren Penko, Cristina Rubino, Anjana Rajendran, Ronan Denyer, Christina B. Jones, Jordan Brocato, Chris Lamb, Cristina Schaurich, Tamara Koren, Lara Boyd, Sarah N. Kraeutner

**Affiliations:** Neuroplasticity, Imagery, and Motor Behaviour Laboratory, Department of Psychology, University of British Columbia, Kelowna, British Columbia, Canada; Graduate Program in Rehabilitation Sciences, University of British Columbia, Vancouver, British Columbia, Canada; Graduate Program in Neuroscience, University of British Columbia, Vancouver, British Columbia, Canada; Graduate Program in Experimental Medicine, University of British Columbia, Vancouver, British Columbia, Canada; Department of Physical Therapy, University of British Columbia, Vancouver, British Columbia, Canada; Djavad Mowafaghian Centre for Brain Health, University of British Columbia, Vancouver, British Columbia, Canada; Department of Psychology, Irving K. Barber Faculty of Arts and Social Sciences, University of British Columbia, Kelowna, Canada; Institute of Neuroscience, Université Catholique de Louvain, Brussels, Belgium

**Author notes:** Address for Correspondence Dr. S. N. Kraeutner, Department of Psychology, the University of British Columbia Okanagan Campus, Rm 204, Arts and Sciences (ASC) 1147 Research Rd, Kelowna, BC Canada, V1V 1V7.

**Keywords:** mental practice, brain injury, neuroimaging, spatial analyses, sex differences

## Abstract

Motor imagery (MI; the mental rehearsal of movement) activates sensorimotor regions, providing the basis for its effectiveness as an intervention for motor recovery after stroke. Yet, the impact of biological sex on MI-related brain activation after stroke is unexplored. Here, we investigated sex-related differences in MI-related brain activation after stroke and explored associations between questionnaire-based MI ability and MI-related brain activation. Thirty-four individuals with chronic stroke performed MI of a complex, upper-limb movement using their paretic arm/hand. Brain activation was captured using functional magnetic resonance imaging (fMRI). Individual differences in MI ability were also assessed prior to the scan using an established MI questionnaire. MI-related brain activation was observed across sensorimotor regions. Yet, group-level contrasts revealed greater activation in sensorimotor regions for females vs. males, while greater activation in the cerebellar and parietal regions was noted in males vs. females. Females also recruited greater ipsilesional regions vs. males. Questionnaire-based MI ability was associated with brain activation localized to occipital regions. Our findings suggest that females and males may respond differently to MI-based tasks. Further, we did not find a relationship between questionnaire-based data and activity in sensorimotor regions. This finding suggests that multiple measures of MI may be needed to assess its impact. Overall, this work informs the use of MI after stroke.

---

Chronic (>6 months post stroke) upper-limb movement impairments experienced by 85% of stroke survivors are related to decreased quality of life (Go et al., 2014). Upper-limb movement impairments following a stroke are linked with changes in brain function in the sensorimotor network (Askim et al., 2009; Dong et al., 2006). Learning is central to restoring function after stroke; practice drives the formation and strengthening of neural pathways that are essential for re-establishing motor control and coordination (Kleim & Jones, 2008). Indeed, physical practice is considered the gold standard to stimulate neuroplasticity. Yet, as current rehabilitation practices do not stimulate sufficient recovery (Langhorne et al., 2011), motor imagery (MI; the mental rehearsal of a movement) has emerged as a promising adjunct to physical practice (Braun et al., 2008; Malouin et al., 2013). MI has been shown to be effective to promote learning in numerous domains (Ruffino et al., 2021; Williams et al., 2015). The basis for the effectiveness of MI is derived from the theory of functional equivalence; MI shares neural processes with physical practice (Hardwick et al., 2018; Hétu et al., 2013, Jeannerod, 1995; Kraeutner et al., 2014; Newell., 1991), driving neuroplasticity necessary for learning and recovery. However, there is evidence of variable response to MI after stroke (Braun et al., 2006; Dickstein & Deutsch., 2007; Ietswart et al., 2011; Sharma et al., 2009), yet this variability has not been extensively investigated.

Past work using functional magnetic resonance imaging (fMRI) to investigate patterns of brain activation during MI in stroke showed that, relative to healthy controls, brain activation was localized to similar regions between groups. However, MI-related activity was noted to be at a lesser degree in individuals with stroke, observed via reduced connectivity between motor and premotor regions, as well as smaller regions of activation (Sharma et al., 2009). Specifically, both healthy controls and individuals after stroke demonstrated enhanced task-specific cortical activation during MI of a finger-thumb opposition task, and this activation was localized to motor and sensorimotor regions (i.e., prefrontal cortex, dorsal premotor cortex, primary motor cortex, cerebellum; Sharma et al., 2009). However, unique to individuals after stroke, the supplementary motor area and cerebellum showed larger regions of activation (Sharma et al., 2009). However, the authors did not examine whether other factors could be driving these differences. Further, while MI ability was assessed (i.e., the capacity to accurately perform MI as assessed by questionnaires), the authors excluded participants with low MI ability from analyses; thus, it is not clear how differences in MI ability might relate to different patterns of brain activation. In work by Kimberley et al. (2006), while healthy individuals primarily activated the contralateral supplementary motor area when performing MI of an upper-limb task, individuals with stroke primarily activated the ipsilateral supplementary motor area. Together, these studies highlight the variability in response to MI after stroke. To begin to understand the source of this variability new data are needed that consider potential sources of differences in response to MI such as sex and MI-ability.

Andrushko et al. (2023) investigated differences in functional brain activation between males and females during a voluntary movement task. This work showed that females have smaller volumes of brain activation and lower patterns of variability across the movement tasks (Andrushko et al., 2023). As it is established that MI-related brain activation overlaps with that of physical practice (Hardwick et al., 2018; Hétu et al., 2013; Jeannerod, 1995; Kraeutner et al., 2014; Newell., 1991), these findings highlight the role that sex might have on shaping brain activation patterns during physical practice, which may translate to MI practice. Given that females experience worse physical function and quality of life outcomes after stroke (Guo et al., 2023; Lai et al., 2005), along with evidence of sex differences in brain activation during physical practice (Andrushko et al., 2023), investigating sex differences in MI-related brain activation after stroke may inform rehabilitation strategies.

Another important factor that has not been explicitly considered is the potential impact of individual differences in MI-ability on resultant brain activation patterns. While MI ability may be reduced depending on location of stroke-related damage, past work investigating brain activation during MI typically screens for MI ability rather than interrogating its impact (Kimberley et al., 2006; McInnes et al., 2015; Oostra et al., 2016; Sharma et al., 2009; Sirigu et al., 1996). Further, the questionnaire-based approach typically used to assess MI ability may be better suited to capture the vividness of one’s MI (Baddeley et al., 2000; Kraeutner et al., 2020; Kraeutner & Scott, *accepted*), and it is unclear whether or how these questionnaires reflect sensorimotor activation during MI after stroke.

Accordingly, the current study examined whether response to MI in individuals with upper-limb hemiparesis after stroke differs across sex. Participants performed MI of a complex upper-limb task while undergoing fMRI. In line with past work (Sharma et al., 2009), we hypothesized that MI-related brain activation would be observed across sensorimotor areas. Further, we expected sex-specific differences, with more focal brain activity in sensorimotor areas for females, compared to more widespread brain activity in visual and sensorimotor areas in males. As a secondary aim, we explored associations between questionnaire-based reports of MI ability and MI-related brain activation after stroke. We expected that greater kinaesthetic MI ability would be related to a higher magnitude of brain activation within sensorimotor areas.

## METHODS

### Participants

Thirty-six adults aged between 30 and 90 years (67.3 ± 13.2), with upper-limb hemiparesis as a result of stroke, were recruited for this study. Two participants were excluded due to technical errors in the scan, and a stroke secondary to a traumatic brain injury, leaving thirty-four participants included in final analysis (12 females; 15 left hemisphere stroke). Sample size was determined via two power analyses conducted using free statistical software (G*Power 3.1). For sex-related analyses, it was determined that 23 total participants would achieve statistical power (Linear multiple regression: fixed model, R^2^ increase: α = .05, β = .1, η_p_^2^ = .34, Critical *F* = 4.32). A large effect size was used based on previous studies examining the difference in brain activation patterns following stroke (Park et al., 2015). We subdivided our participants so that 12 males who were age-matched to the 12 females in our sample were used to test our questions surrounding the impact of sex on MI (mean age of females and males 69.6 ± *12.5*). For MI ability-related analyses, it was determined that 25 participants would achieve statistical power (Linear multiple regression: fixed model, R^2^ increase: α = .05, β = .2, η^2^ = 8.75, Critical *F* = 4.28). A large effect size was used based on previous studies examining the effects of MI-ability on brain activation patterns (Lee et al., 2019). Thus, our overall sample size was powered to conduct this analysis. Ethical approval was granted by the institution’s Research Ethics Board. All participants provided written, informed consent and were oriented to the experimental task by the experimenter. Note that the data reported here represents baseline data from work reported in Magnuson et al., *in review*, where participants engaged in a longitudinal study with the goal to determine the effects of non-invasive brain stimulation on task-related brain activity after stroke.

Before beginning the experiment, all participants completed the Kinesthetic and Visual Imagery Questionnaire (KVIQ; Malouin et al., 2007) to characterize their MI ability in two domains: kinesthetic (“the feelings and sensations experienced if you were actually producing the movement”), and visual (“when you watch yourself performing the movement from an outside point of view or third-person perspective”). Participants also completed the Upper-Extremity Fugl-Meyer Assessment (FMA-UE) to assess the degree of motor impairment (Fugl-Meyer., 1975; Singer & Garcia., 2017).

### Data acquisition

Participants were placed in an MRI scanner (Phillips 3.0T whole body MRI scanner with an 8-channel sensitivity encoding head coil (SENSE factor = 2.4) and parallel imaging) and were given sound-cancelling headphones to reduce distractions as well as to provide communication through. Structural (High-resolution T_1_ scan: TR = 1900 ms, FOV = 256 mm, 160 slices, 1 mm thickness, 3.2 min, and T2-weighted anatomical image: TR = 4200 ms, FOV = 256 mm, 184 slices, 1 mm thickness, 6 min, for lesion identification and co-registration purposes) and functional (blood-oxygen level dependent (BOLD) scans with a single shot EPI sequence (TR = 2000 ms, TE = 30 ms, FOV = 240 mm, 105 slices, 2.5 mm thickness, 6 min/scan) images were obtained during the MRI session.

The fMRI experiment included a single run involving two conditions: MI and rest. Blocks began and ended with a rest block (30-second) and included three 30-second MI task blocks alternating with rest. Blocks began with a visual cue (“Imagine” and “Rest” on a screen situated above the participant using the Presentation software package, synchronized with the Phillips MRI system, followed by a 30-second response period. When the screen presented “Imagine” participants performed an MI task which involved imagining opening a door using their affected hand (without physically moving, therefore not relying on one’s physical ability to perform the movement); a task with a high degree of familiarity to the participant has been shown to make brain activation more focal and lateralized (Kraeutner et al., 2018). Participants laid on their back in the scanner, with their hands resting comfortably in their lap. Immediately following the MI task, and while still lying inside the scanner, participants were asked to rate their engagement and quality of imagery on a scale of 1 (not engaged; poor quality) to 5 (extremely engaged; excellent quality).

### Experimental protocol

Participants underwent two visits to the Brain Behaviour Laboratory and Charles E. Fipke Integrated Neuroimaging Suite over two consecutive days, at least 24 hours apart, at the University of British Columbia. The first visit involved baseline assessments (FMA-UE, KVIQ). During the second visit, participants completed the MRI session. Prior to the MRI session participants were oriented to the task and type of imagery to be performed in the scanner (kinesthetic; first person; Moran et al., 2012; Stinear et al., 2007).

### Data analysis

#### Pre-processing

MRI pre-processing was carried out using fMRIprep version 23.0.0 (Smith., 2002). Anatomical MRI was used for co-registration with functional data. Anatomical T_1_ images were corrected for intensity non-uniformity using N4BiasFieldCorrection and skull-stripped using antsBrainExtraction using the OASIS template. Spatial normalization was performed through nonlinear registration with ANT’s antsRegistration tool. Brain surfaces were reconstructed using recon-all from Freesurfer version 7.4.1. Pre-processing of functional images included motion correction using FSL’s MCFLIRT followed by co-registration to the anatomical T_1_ using FreeSurfer’s bbregister and 6 degrees of freedom. ANT’s antsApplyTransforms was used to apply Lanczos interpolation to the BOLD-to-T_1_ transformation and T_1_-to-MNI warp. ICA-based Automatic Removal of Motion Artifacts (ICA-AROMA) was used to generate non-aggressively denoised data. Brains were flipped along the x-axis prior to statistical analyses, such that all lesions were represented in the right hemisphere of the brain (henceforth we refer to the right hemisphere as the ipsilesional hemisphere).

#### Statistical analysis

Individual statistical activation maps were calculated for the run using GLM with FEAT then second-level averaged across runs to produce task activation maps for each participant, for group level analyses. To determine whether, and to what extent MI-related activity is modulated by sex differences, separate higher-level (group) analyses were carried out across all participants and disaggregated by sex using Fixed Effects (FMRIB’s Local Analysis of Fixed Effects). We performed a contrast analysis to assess areas where males exhibit significantly greater brain activation during MI than females using Fixed Effects (coded +1 for males; −1 for females). Conversely, a second contrast was conducted to identify regions where females showed greater activation relative to males using Fixed Effects (coded −1 for males, +1 for females). All analyses were thresholded using clusters determined by Z > 3.1, and a corrected cluster significance threshold (p = 0.05).

## RESULTS

Demographic data for each participant is shown in Table 1 (Kinesthetic KVIQ: *M =* 17.9 ± 5.3; Visual KVIQ: *M =* 20.6 ± *SD =* 4.9; MI quality: *M =* 3.7 ± 1.2; MI engagement: *M =* 3.9 ± 1.2).

**Table 1.**
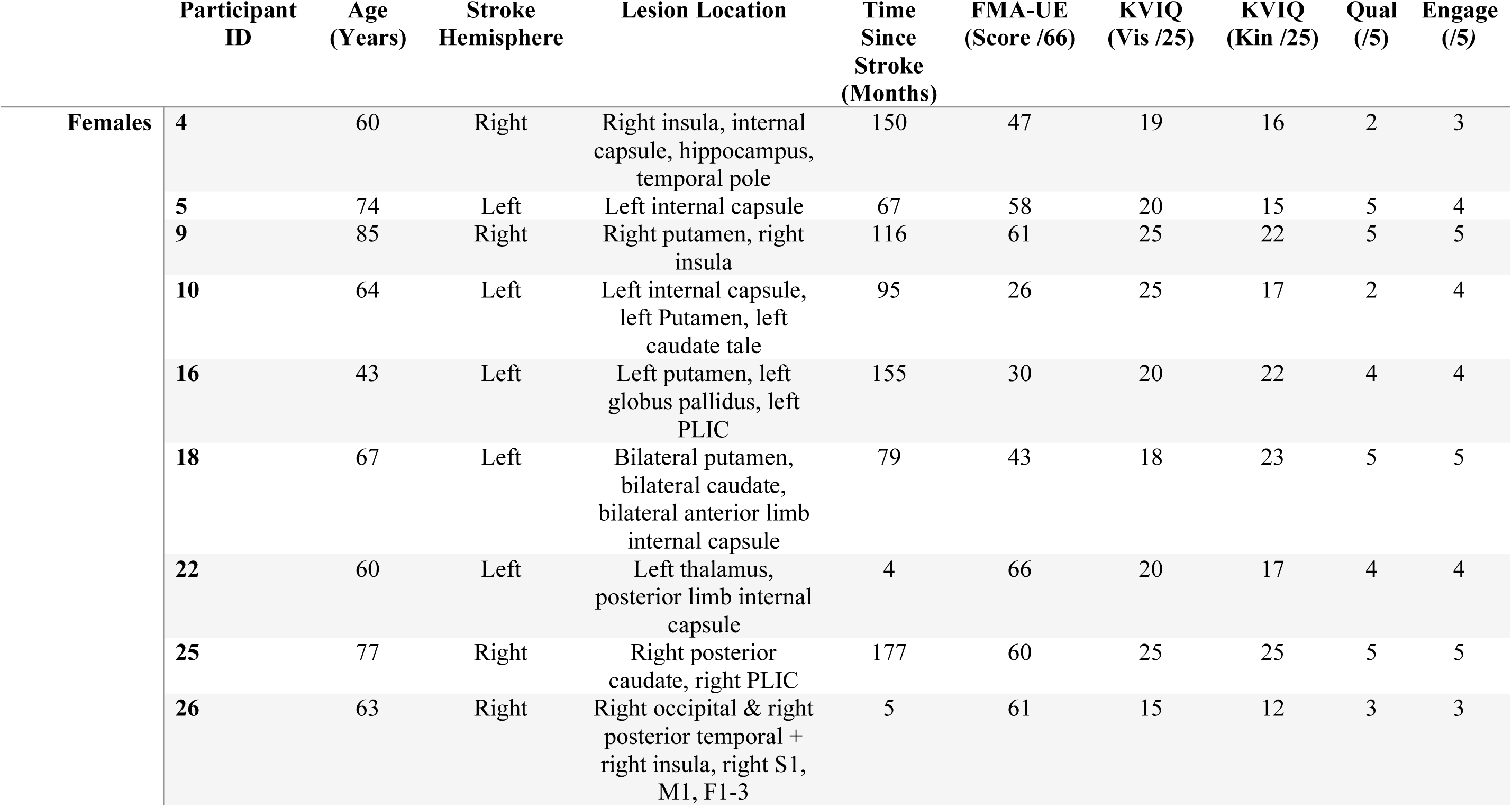
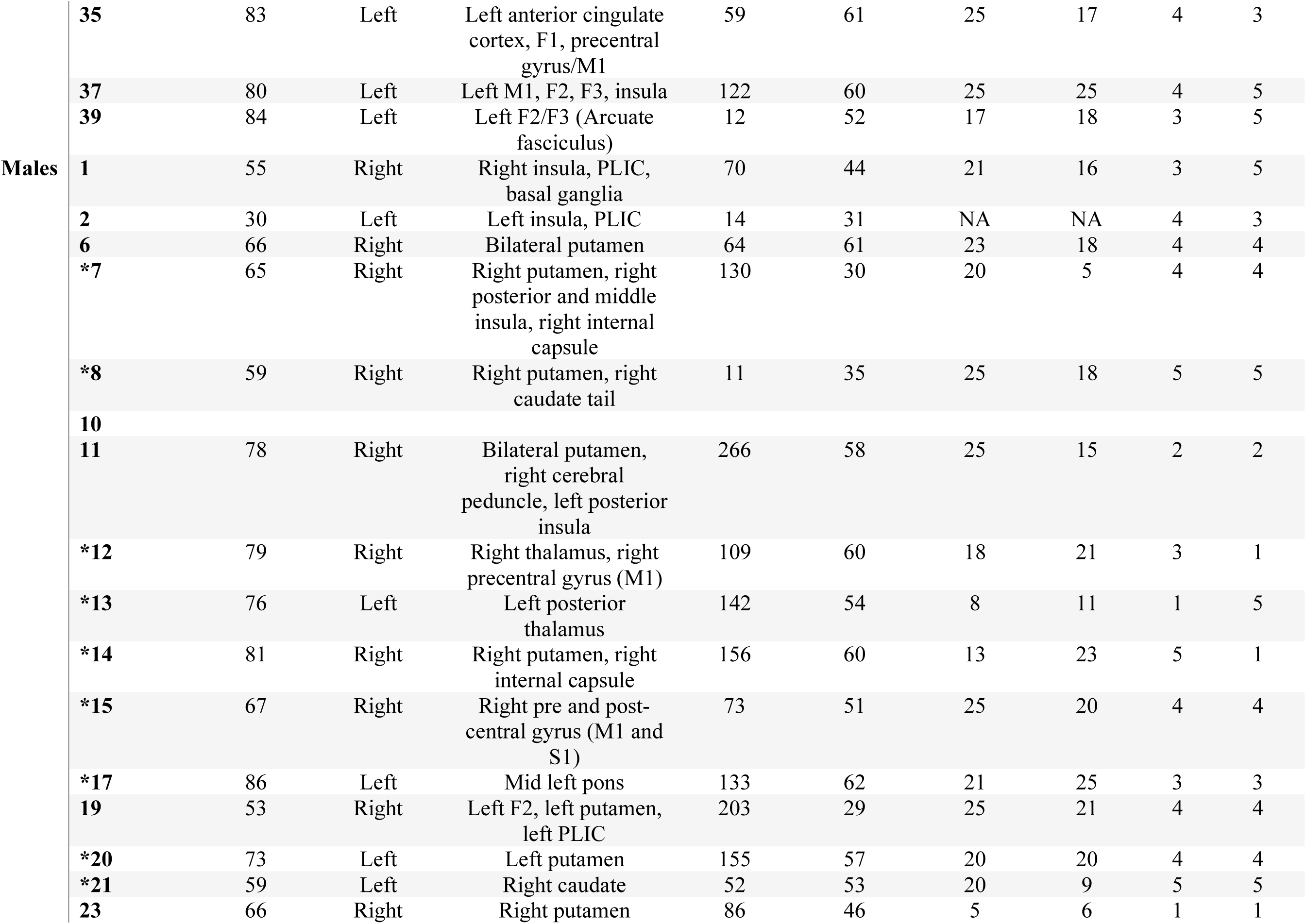
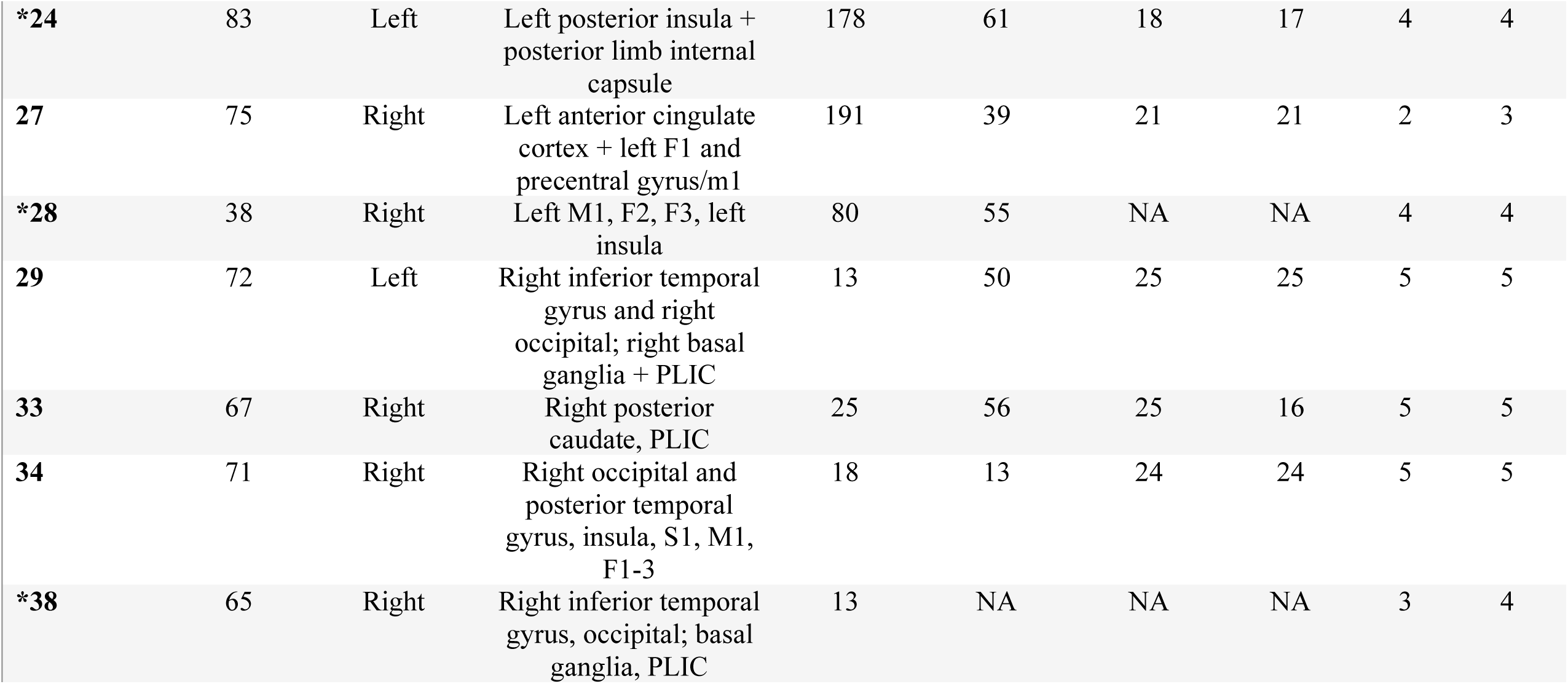
Participant demographics. Demographics include age, sex, Upper Extremity Fugl Meyer score (FMA-UE), both visual and kinesthetic domains of the Kinesthetic and Visual Imagery Questionnaire (KVIQ), time since stroke (calculated in months, lesion location,) and MI quality (qual) and engagement (engage) ratings.(Abbreviations: PLIC: posterior limb of internal capsule; S1: primary somatosensory cortex; M1: primary motor cortex; F1: Brodmann area 4; F2: caudal part of superior area 6 (part of premotor cortex); F3: frontal mirror neuron system). * Denotes which males were included in the sex differences analysis.

### Motor imagery-related brain activation

Group-level brain activation (regardless of sex) during MI is reported in Table 2. Overall, MI-related brain activation was observed in sensorimotor regions (including bilateral precentral gyri), cerebellar regions (including ipsilesional cerebellum crus I; Figure 2).

**Figure 1.**
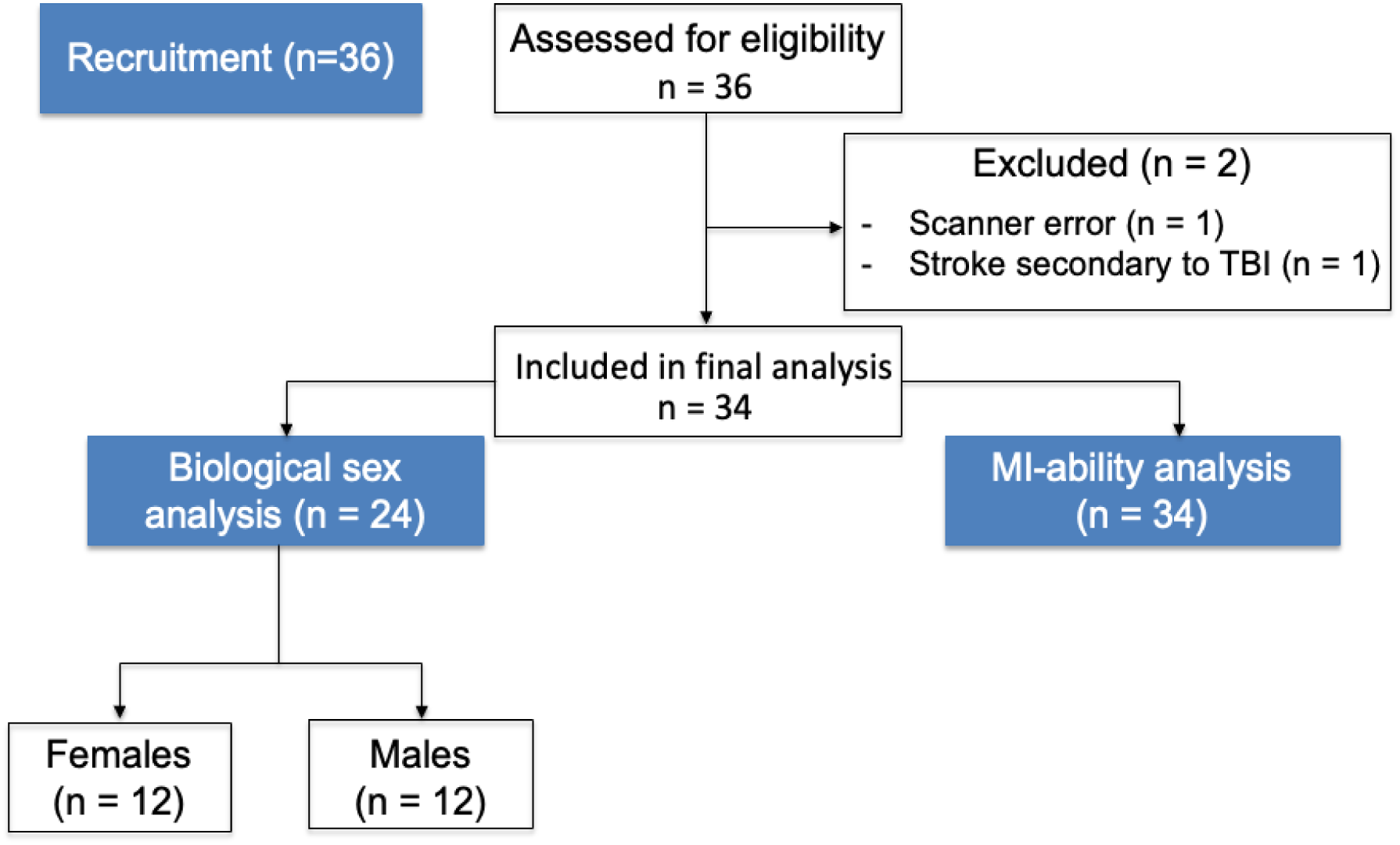
Flowchart of participant inclusion and analysis. A total of 36 participants were recruited for the study. Two participants were excluded: one due to scanner errors and another due to a stroke secondary to traumatic brain injury (TBI), leaving 34 participants for the final analysis. All 34 participants were included in the motor imagery ability analysis. For the sex-based subgroup analysis, 12 males were age-matched to 12 females (n=24).

**Figure 2.**
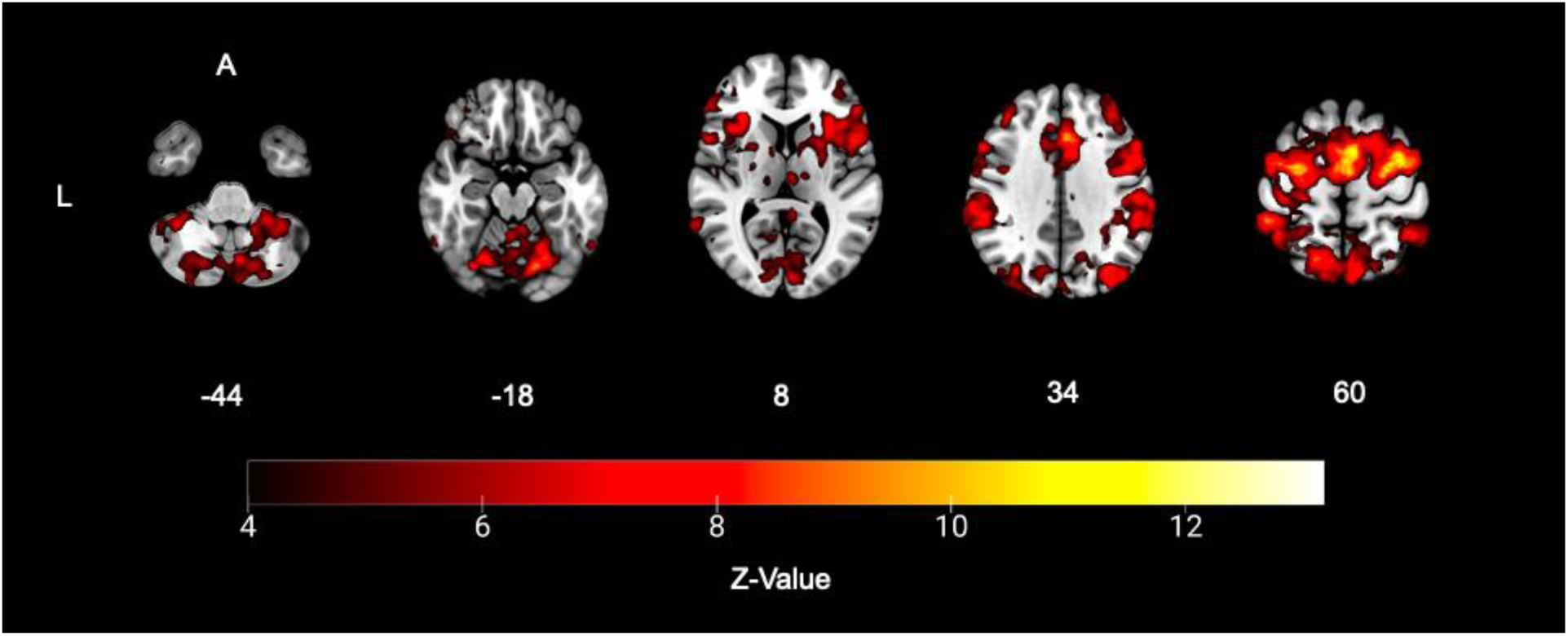
Second-level activation maps showing overall brain activation during MI for all participants (group-mean). Warm colors indicate greater magnitude of activation in overlayed regions. The color bar represents z-max values. Activation was localized to ipsilesional cerebellum crus1; contralesional fusiform gyrus, inferior temporal gyrus; and bilateral superior frontal gyrus, and precentral gyrus.

**Table 2.** MNI coordinates of local maxima of clusters of significant activation for MI-related brain activation across all participants.

| Hemisphere | *Anatomical location* | MNI coordinates (mm) |  |  | *Z value* |
| --- | --- | --- | --- | --- | --- |
|  |  | *x* | *y* | *z* |  |
| Ipsilesional | Superior frontal gyrus | 26 | -8 | 64 | 13.6 |
|  | Precentral gyrus | 26 | -18 | 74 | 12.7 |
|  | Cerebellum crus I | 38 | -54 | -28 | 12.3 |
| Contralesional | Superior frontal gyrus | -32 | -2 | 68 | 13.1 |
|  | Precentral gyrus | -54 | 4 | 46 | 12.2 |
|  | Fusiform gyrus | -36 | -4 | -48 | 5.61 |
|  | Inferior temporal gyrus | -54 | -4 | -44 | 4.86 |

MI-related activation across sex is reported in Table 3 and shown separately in Figure 3. Between-group contrasts conducted to assess MI-related brain activation between sex (females vs. males) indicated that overall, greater activation across ipsilesional areas was observed for females (9 regions) vs. males (4 regions). Greater activation for females vs. males was observed across sensorimotor regions (encompassing contralesional precentral gyrus and inferior parietal lobule, and ipsilesional postcentral and supramarginal gyri), frontal regions (including ipsilesional middle orbitofrontal gyrus and contralesional middle frontal gyrus), occipital regions (including ipsilesional middle occipital gyrus, contralesional superior occipital and lingual gyri), and temporal regions (including bilateral middle and inferior temporal gyri; Figure 4). Greater activation for males vs. females was observed across bilateral cerebellar regions and bilateral parietal areas (encompassing ipsilesional angular gyrus, and contralesional superior and inferior parietal lobules; Figure 4).

**Figure 3.**
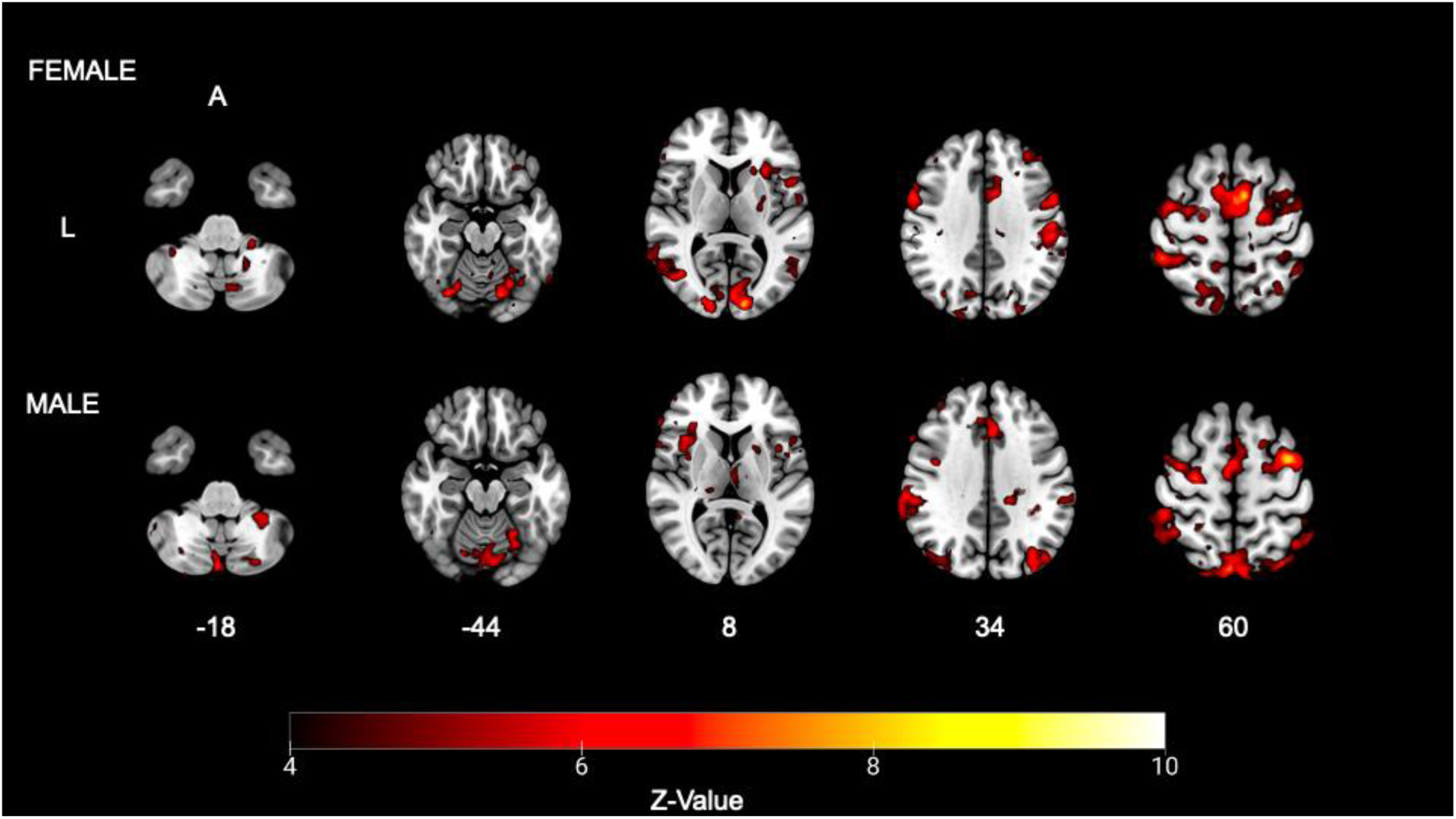
Higher-level activation maps showing MI-related brain activation for females (**top)** and males **(bottom);** the color bar represents *Z*-max values. Warm colors indicate a greater magnitude of activation in the overlayed regions. For females, activation was localized to the ipsilesional supramarginal gyrus, inferior parietal lobule, inferior orbitofrontal gyrus, frontal inferior operculum, middle orbitofrontal gyrus, and bilateral supplementary motor areas, and precentral gyrus. For males, activation was localized to ipsilesional cerebellum crus1, inferior temporal gyrus, calcarine, insula, caudate; contralesional cerebellum VIII, middle frontal gyrus, inferior frontal gyrus cuneus; and bilateral precentral gyrus.

**Figure 4.**
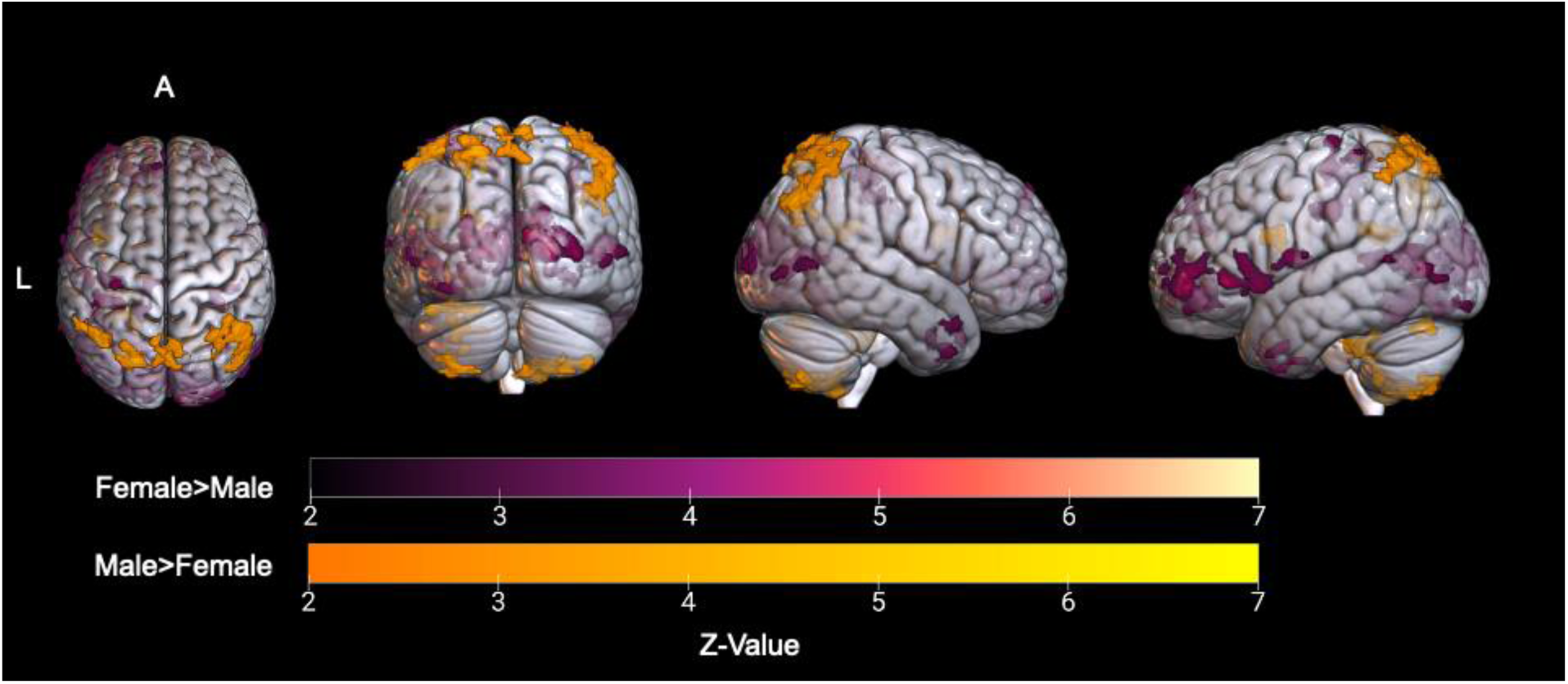
Sex-related differences on resultant MI-based brain activation; the color bar represents the *Z*-max value. Purple colors indicate regions that had greater magnitude of activation in females when compared to males. Yellow colors indicate regions that had greater magnitude of activation in males when compared to females. Activated voxel clusters for females vs. males were localized to ipsilesional middle occipital gyrus, supramarginal gyrus, middle orbitofrontal gyrus; contralesional precentral gyrus, precuneus, paracentral lobule, rolandic operculum, inferior parietal lobule; and bilateral cuneus, superior occipital gyrus, lingual gyrus, postcentral gyrus, and inferior temporal gyrus. Activated voxel clusters for males vs. females were localized to ipsilesional angular gyrus; contralesional superior parietal lobule, cerebellum crus II, inferior parietal lobule, cuneus, inferior frontal operculum; and bilateral precuneus, cerebellum VIIb, and cerebellum VIII.

**Table 3.**
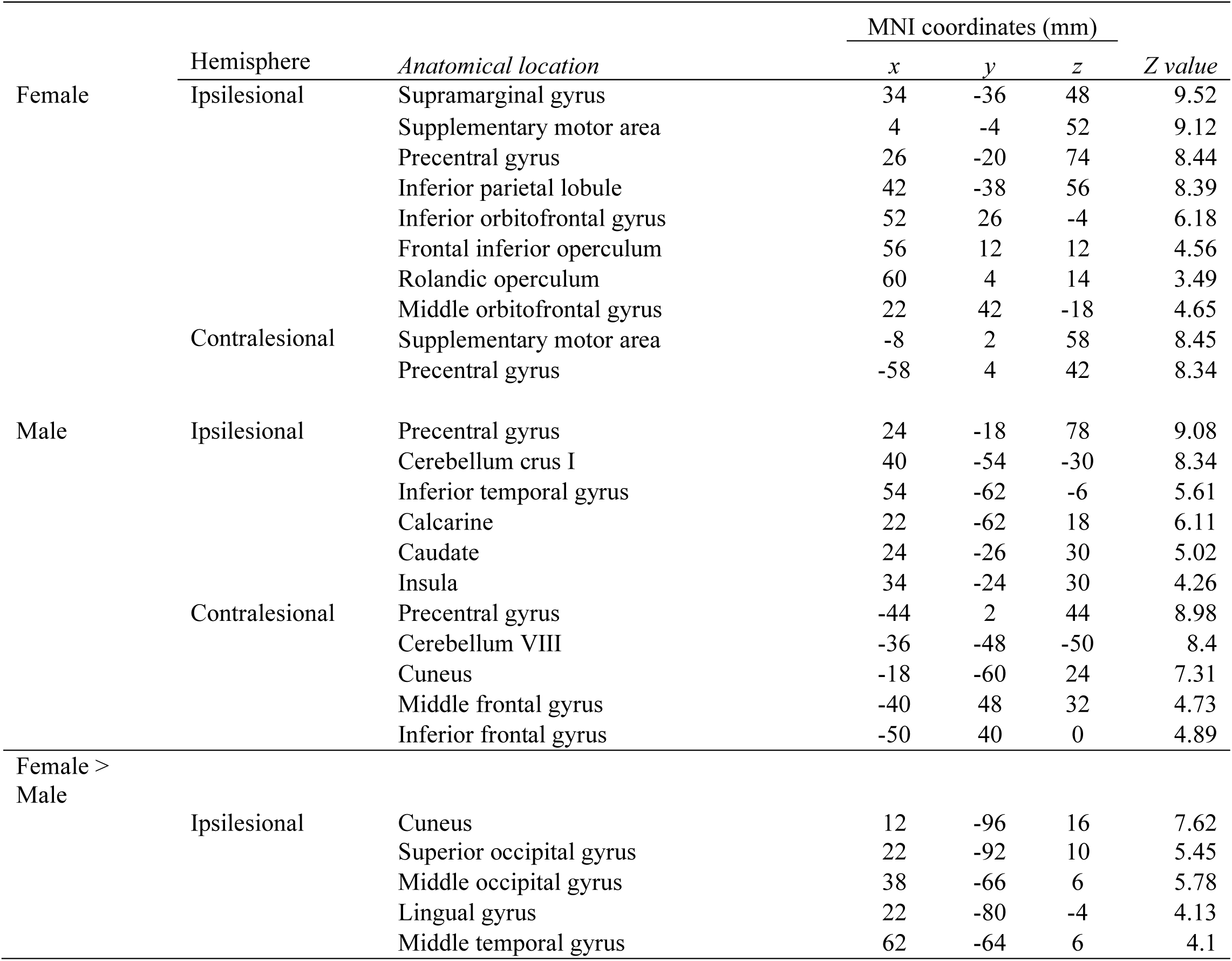
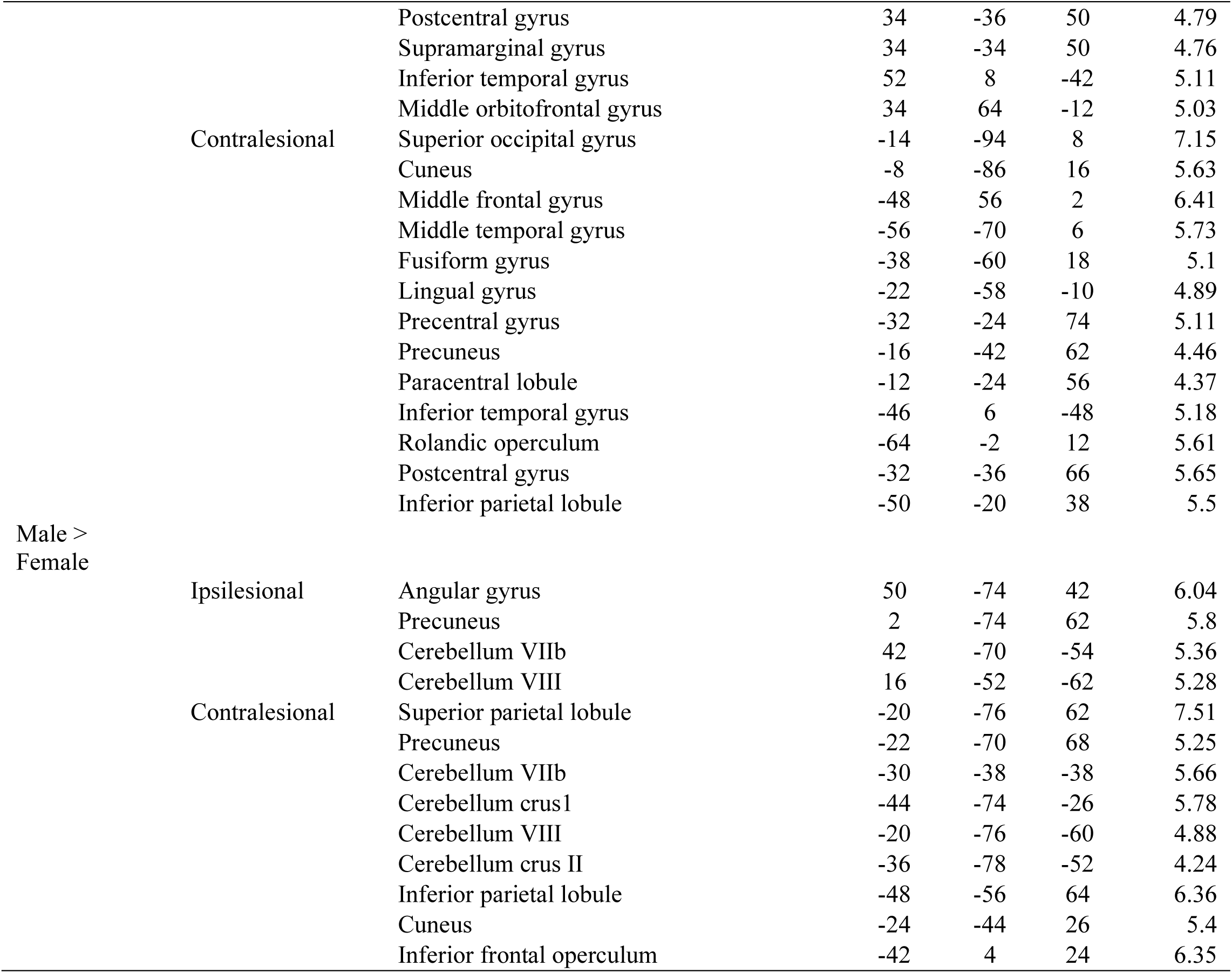
MNI coordinates of local maxima of clusters of significant activation for MI-related brain activation across sex and resulting from comparisons conducted to assess sex-related differences in motor imagery-related brain activation.

### MI ability and brain activation during MI

Results from our exploratory analysis testing the association between MI-ability and brain activation during MI after stroke are reported in Table 4. Kinaesthetic MI ability was positively associated with activation across bilateral visual regions, including ipsilesional cuneus and superior occipital gyrus, and contralesional middle occipital gyrus, and calcarine (Figure 5).

**Figure 5.**
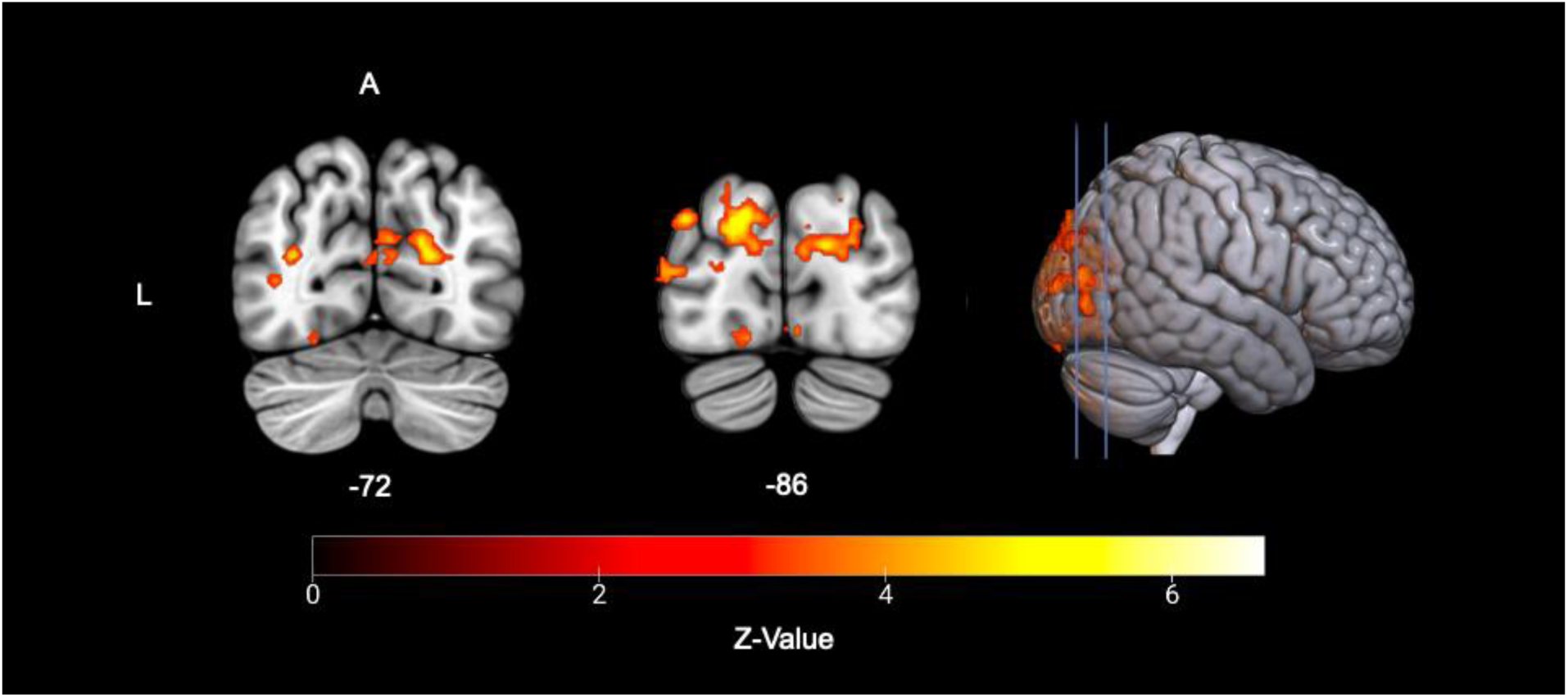
Association between kinesthetic KVIQ score and resultant MI-based brain activation after stroke, where the color bar represents the *Z*-max value. Warn colors indicate regions of activation. Activated voxel clusters are localized to the ipsilesional cuneus and superior occipital gyrus; contralateral middle occipital gyrus, and calcarine.

**Table 4.** MNI coordinates of local maxima of clusters of significant activation for MI-related brain activation associated with kinesthetic scores on the KVIQ.

| Hemisphere | Anatomical location | MNI coordinates (mm) |  |  | Z value |
| --- | --- | --- | --- | --- | --- |
|  |  | x | y | z |  |
| Ipsilesional | Superior occipital gyrus | 12 | -96 | 18 | 8.67 |
|  | Cuneus | 12 | -102 | 12 | 7.22 |
| Contralesional | Calcarine | -2 | -90 | -12 | 10.8 |
|  | Middle occipital gyrus | -12 | -104 | 6 | 6.83 |

## DISCUSSION

In the current study, we assessed sex-related differences in brain activation during MI after stroke. Consistent with prior work, MI-related brain activation overall was observed in sensorimotor regions, including bilateral precentral gyri. In line with our hypotheses, we observed different patterns of activation in females vs. males, with females showing greater recruitment across cortical sensorimotor regions and bilateral frontal and occipital areas. In contrast, males showed greater activation across cerebellar and parietal regions. Kinesthetic MI ability was positively associated with activation in regions implicated in visual, but not sensorimotor processing, suggesting that perceived MI-ability may not be associated with sensorimotor activation after stroke. Below we discuss our findings in the context of prior work examining MI after stroke and neural processes underlying motor tasks.

Our finding that females activated sensorimotor regions to a greater degree compared to males suggests that females may be well-suited for MI-based interventions. As sensorimotor activation is necessary to facilitate motor learning and recovery after stroke (Edwards et al., 2019; Ghilardi et al., 2000; Kleim & Jones, 2008), the effectiveness of MI in facilitating recovery after stroke likely depends on engagement of these regions. While it is well documented that MI recruits sensorimotor regions in healthy individuals (e.gs., Hardwick et al., 2018; Hétu et al., 2013), our findings thus suggest that MI may preferentially drive activation of these areas in females, potentially leading to more effective motor recovery relative to males.

Emerging evidence supports sex-related differences in brain function during motor tasks (Andrushko et al., 2023), with males exhibiting more widespread activation, involving visual and sensorimotor areas, which may reflect a greater reliance on visuospatial processing (Andrushko et al., 2023; Bianco et al., 2020). In contrast, females generally recruited a greater number of ipsilesional areas (9) relative to males (4), as well as contralesional precentral gyrus. Ipsilesional patterns of activation encompassing core motor regions are crucial for motor recovery as they are indicative of the brain’s efforts to reorganize and compensate for damage (Kleim & Jones, 2008; Serrien et al. 2004). Given that females were shown to recruit activity across both ipsilesional and contralesional areas during MI vs. males in the current study, whether this translates to improved recovery warrants further investigation.

Taking our findings together with the under-recruitment of females in stroke rehabilitation research (Mehrabi et al., 2024), our data may help explain the variable response to MI after stroke noted in past studies, which showed effect sizes ranging from negligible to moderate (Braun et al., 2006; Dickstein & Deutsch., 2007; Ietswart et al., 2011; Sharma et al., 2009). A systematic review of 1209 randomized controlled trials for upper limb stroke rehabilitation found that only 38.8% of their participants were females (Mehrabi et al., 2024), despite females accounting for 56% of the stroke population (Yoon & Bushnell., 2023). This disparity (i.e., proportion of females vs. males) may confound the variability of MI response observed across past work, underscoring the need for the effectiveness of MI for motor recovery after stroke to be tested separately for females vs. males in future work.

In our study, males uniquely activated cerebellar regions, suggesting a greater reliance on processes attributed to these regions to perform MI after stroke. Past work (collapsed across sex) demonstrates that cerebellar activation during motor imagery is modulated by learning (Lacourse et al., 2005). Discrepancies in predicted vs. actual consequences of a movement (via feedback), resulting in large changes to the motor program, modulates cerebellar activation (e.g., Dayan & Cohen, 2011; Lacourse, Turner, Randolph-Orr, Schandler, & Cohen, 2004; Seitz et al., 1994). Further, the cortico-pontine-cerebellar tract links the upper part of the posterior cerebellum to the supplementary motor area and the premotor cortex (Middleton and Strick, 1994), relying on information important for movement coordination and inhibition (Henschke & Pakan., 2023; Rao et al., 1997). Males also demonstrated greater recruitment of posterior parietal regions critical to MI performance (Davis et al., 2024; Kraeutner et al., 2019; Sirigu et al., 1996; Solomon et al., 2022) and MI-based learning (Frank et al., 2023; Kraeutner et al., 2016). For instance, performance on an implicit MI task (requiring individuals to determine if a hand presented on a screen is a left or right hand; termed, the hand laterality judgment task), was disrupted when transcranial magnetic stimulation was delivered over the inferior parietal lobe. While sex was not tested in this work, other work has also shown males more quickly responded to hands presented in the palm view, while females were faster to respond to hands presented in the dorsal view (Conson et al., 2020). In light of the current findings, males and females may use distinct MI strategies after stroke, likely related to how a motor image is transformed in the mind.

It is important to consider why kinaesthetic MI ability was related to occipital activation implicated in vividness of the generated motor image (Oostra et al., 2016) yet was unrelated to activation across sensorimotor regions as hypothesized. It is possible that participants were unable to perform kinaesthetic MI. However, males and females demonstrated MI-related brain activation in sensorimotor regions typically associated with kinesthetic components of MI (Hétu et al., 2013; Oostra et al., 2016; Ptak et al., 2017). Thus, it may well be that KVIQ scores may not reflect activation across sensorimotor areas and instead capture a different aspect of MI. Past work indicates the KVIQ more greatly captures one’s ability to generate motor images (Kraeutner et al., 2020), while other assessments may be more suited to assessing timing and/or transformation components that are more closely associated with sensorimotor processes. Thus, additional measures may be needed to provide a comprehensive assessment of MI ability and/or to ensure its effective prescription for motor recovery after stroke (Gulliot & Collet., 2005; Kraeutner et al. 2020; Kraeutner & Scott, *accepted*).

Here, we investigated whether response to MI in individuals with upper-limb hemiparesis after stroke differs across sex and explored associations between MI-ability and brain activation during MI. Females showed greater activation across sensorimotor regions as well as bilateral frontal and occipital areas relative to males. Our findings suggest that males and females may use different MI strategies after stroke, sub-served by distinct neural processes. Further, we showed that questionnaire-based kinesthetic MI ability reflected activation in occipital (vs. sensorimotor) regions associated with the generation of a motor image, supporting the need to use multiple assessments of MI ability prior to its prescription after stroke. Overall, we recommend that future work investigating MI-based interventions for upper limb recovery after stroke assess effectiveness separately for males and females.

## Author contributions

SMK: data curation, formal analysis, visualization, writing – original draft and review and editing.

JM: data curation, formal analysis, writing – review and editing.

LP: investigation, data curation, methodology, writing – review and editing.

CR: investigation, formal analysis, writing – review and editing.

AR and RD: investigation, data curation, writing – review and editing.

CJ: investigation, writing – review and editing.

JB: project administration, investigation, writing – review and editing.

CL, CS: investigation, writing – review and editing.

TK: project administration, investigation, writing – review and editing.

LB: conceptualization, methodology, writing – original draft and review and editing, funding acquisition, supervision, project administration.

SNK: conceptualization, methodology, writing – original draft and review and editing, funding acquisition, supervision, project administration.

## Statements and declarations

*Ethical considerations:* All procedures were approved by the institutional clinical research ethics board (H24-01280).

*Consent to participate:* Participants gave written informed consent prior to testing

*Consent for publication:* Not applicable

*Declaration of conflicting interest:* The author(s) declared no potential conflicts of interest with respect to the research, authorship, and/or publication of this article

## Acknowledgements

The authors disclosed receipt of the following financial support for the research, authorship, and/or publication of this article: Funding for this work was provided by the Lotte & John Hecht Memorial Foundation made to LB and SNK (ID: 4836). SNK was supported by an early career award through Michael Smith Health Research BC (MSHRBC). JM was supported by a trainee awards from MSHRBC. SMK and CL were supported by CIHR Canada Graduate Scholarships Masters awards. CL was also supported by the Djavad Mowafaghian Centre for Brain Health General Award, the President’s Academic Excellence Initiative PhD award, and the University of British Columbia Rehabilitation Medicine Alumni Jane Hudson Scholarship. RD was supported by the Canadian Partnership for Stroke Recovery.

